# CREB non-autonomously regulates Reproductive Aging through Hedgehog/Patched Signalling

**DOI:** 10.1101/2020.03.02.972018

**Authors:** Nicole M. Templeman, Vanessa Cota, William Keyes, Rachel Kaletsky, Coleen T. Murphy

## Abstract

Evolutionarily conserved signaling pathways are crucial for adjusting growth, reproduction, and cell maintenance in response to altered environmental conditions or energy balance. However, we have an incomplete understanding of the signaling networks and mechanistic changes that coordinate physiological changes across tissues. We found that loss of the cAMP response element-binding protein (CREB) transcription factor significantly slows *Caenorhabditis elegans*’ reproductive decline, an early hallmark of aging in many animals. Our results indicate that CREB acts downstream of the transforming growth factor beta (TGF-β) Sma/Mab pathway in the hypodermis to control reproductive aging, and that it does so by regulating a Hedgehog-related signaling factor, WRT-10. Overexpression of hypodermal *wrt-10* is sufficient to delay reproductive decline and oocyte quality deterioration, potentially acting via Patched-related receptors in the germline. This TGF-β/CREB/WRT-10 signaling axis allows a key metabolic tissue to communicate with the reproductive system to regulate oocyte quality and the rate of reproductive decline.

**Highlights:**

- The transcription factor CREB regulates oocyte quality and reproductive decline
- Hypodermal CREB downstream of TGF-β Sma/Mab signaling controls reproductive aging
- CREB targets in the hypodermis include a Hedgehog-related signaling factor, *wrt-10*
- WRT-10 regulates reproductive aging, potentially via germline Patched receptors

**eTOC Blurb:** Templeman *et al*. describe how CREB, a highly conserved transcription factor, is a central regulator of age-dependent reproductive decline. CREB acts downstream of the TGF-β Sma/Mab pathway in the hypodermis, a key *Caenorhabditis elegans* metabolic tissue, to control oocyte quality maintenance and the rate of reproductive decline. Hypodermal CREB affects the expression of *wrt-10*, which encodes a Hedgehog-related signaling factor. The authors show that wrt-10 regulates reproductive aging, potentially via Patched-related receptors in the germline.

## Introduction

Major signaling pathways that regulate growth and metabolism often exert controls over other key biological functions, such as somatic tissue maintenance, lifespan, and reproduction. Nutrient abundance activates signaling pathways that promote energy-expensive processes such as growth and reproduction; conversely, nutrient depletion leads to molecular changes that instead favor cell maintenance and stress resistance, and these changes can ultimately prolong reproductive capacity and extend lifespan (Templeman and Murphy, 2018). Importantly, the signaling pathways and molecular mechanisms governing these processes show a high degree of evolutionary conservation.

A decline in female reproductive capacity, or ‘reproductive aging,’ is one of the earliest hallmarks of age-related deterioration in humans. The increasing occurrences of birth defects, infertility, miscarriage, and unsuccessful pregnancy outcomes that characterize the initial stages of reproductive decline emerge in the decade before fertility ends, and are likely due to an age-related deterioration in oocyte quality (Armstrong, 2001; te Velde and Pearson, 2002). Interestingly, there are close ties between reproduction and processes in the rest of the body, including metabolic homeostasis and somatic tissue maintenance. For instance, in humans, prolonged female fertility is associated with an increase in life expectancy (Gagnon, 2015; Perls et al., 1997). Experiments in model organisms such as *Caenorhabditis elegans* have shown that the germline is both responsive to and responsible for signals that coordinate germline and somatic tissue maintenance, nutrient stores, and reproduction (Hsin and Kenyon, 1999; Luo et al., 2010; Tang and Han, 2017). Delineating the signals that direct these inter-tissue relationships might not only characterize reproductive aging, but also uncover mechanisms that control systemic age-related deterioration.

The nematode *C. elegans* has proven to be a useful model to study reproductive aging. Similar to humans, the period of adulthood during which *C. elegans* hermaphrodites can produce progeny lasts for only one-third to one-half of their lifespan, and a deterioration in oocyte quality underlies reproductive decline (Hughes et al., 2007; Luo et al., 2010; 2009). Moreover, many age-dependent transcriptional changes in oocytes are conserved from *C. elegans* to mammals (Hamatani et al., 2004; Luo et al., 2010; Steuerwald et al., 2007; Templeman et al., 2018).

Highly conserved signaling pathways are integral for determining the rate of reproductive decline. For instance, insulin/insulin-like growth factor 1 (IGF-1) signaling has pronounced effects on aging; loss of function of the *C. elegans* insulin/IGF 1 signaling receptor dramatically extends lifespan (Kenyon et al., 1993) and delays reproductive senescence (Gems et al., 1998; Hughes et al., 2007; Luo et al., 2010; Templeman et al., 2018). Transforming Growth Factor β (TGF-β) signaling, another conserved signaling network that coordinates many aspects of development, cell function, and survival, also regulates aging phenotypes (Savage-Dunn and Padgett, 2017; Shaw et al., 2007). Interestingly, the Sma/Mab branch of *C. elegans* TGF-β signaling (homologous to the mammalian Bone Morphogenetic Protein or BMP subfamily of TGF-β ligands and signal transducers; Savage-Dunn and Padgett, 2017), has only minor effects on lifespan while having pronounced effects on growth (Savage-Dunn et al., 2003) and reproductive aging (Luo et al., 2009). Moreover, TGF-β Sma/Mab signaling affects oocyte and germline quality cell non-autonomously, by acting in the hypodermis (Luo et al., 2010), a *C. elegans* epithelial tissue involved in regulating growth (Wang et al., 2002), development and molting (Altun and Hall, 2009), and metabolic processes (Kaletsky et al., 2018). However, it was not yet known how TGF-β Sma/Mab signaling in the hypodermis leads to reproductive changes. Filling in these knowledge gaps and uncovering signaling components involved in inter-tissue communication is essential to fully understand the processes governing reproductive aging.

In a pilot screen for reproductive aging regulators, we found that loss of function of the *C. elegans* cAMP response element-binding protein (CREB) extended reproductive span (S. Luo, W. Keyes, & CT Murphy, unpublished). CREB is a transcription factor that regulates a wide range of functions, including metabolic processes, cellular respiration, cell survival and proliferation, long-term memory, and immune function (Altarejos and Montminy, 2011; Mayr and Montminy, 2001). CREB’s transcriptional activity is responsive to a variety of upstream cues, and is further altered by cellular cofactors or transcription coactivators (Altarejos and Montminy, 2011; Kimura et al., 2002; Mayr and Montminy, 2001). Similar to mammals, CREB is required for longterm associative memory in *C. elegans* (Kauffman et al., 2010), and also affects *C. elegans* growth, development and metabolic processes (Lakhina et al., 2015), pointing to a potential role in regulating aging phenotypes. Although our lab and others showed that severe loss-of-function or null mutation of *crh-1*, the gene encoding the *C. elegans* homolog of mammalian CREB, does not extend lifespan (Chen et al., 2016; Lakhina et al., 2015), its role in reproductive aging had not been previously described.

In this study, we have delineated a new hypodermis-to-oocyte signaling axis in which oocyte quality and age-dependent reproductive capacity are regulated in response to signals derived from a key metabolic tissue. CREB appears to be a lynchpin for regulating reproductive aging in *C. elegans:* it acts downstream of TGF-β Sma/Mab signaling in the hypodermis to control the rate of reproductive decline, and it does so at least in part by regulating the hypodermal mRNA levels of a Hedgehog-related signaling factor, *wrt-10.* We show that hypodermal overexpression of *wrt-10* promotes oocyte quality maintenance and slows reproductive decline, and that the presence of Patched-related receptors in the germline is required for the full extent of these effects. This network of intra- and inter-tissue lines of communication between evolutionarily conserved signaling pathways play important roles in regulating age-related reproductive decline.

## Results

### Loss of CREB signaling delays age-dependent reproductive decline

CREB is a ubiquitously expressed transcription factor that regulates a wide suite of conserved biological functions. In *C. elegans*, CREB is involved in the regulation of growth, development, and metabolic processes (Lakhina et al., 2015; Figures S1A, S1B), suggesting that it could affect age-dependent physiological decline. In the course of determining that CREB does not regulate longevity (Lakhina et al., 2015), we found that it does control the rate of reproductive aging (Figure 1A); simultaneously, *crh-1* emerged as a candidate reproductive aging regulator from a genetic screen in our lab (unpublished data). Using two *C. elegans* strains with different *crh-1* loss-of-function or null alleles, we found that whole-body loss of *crh-1* leads to a significant extension of the mated reproductive span *(i.e.*, the period of adulthood during which mated hermaphrodites are capable of progeny production, without being limited by sperm quantity), compared to wild-type N2 animals (Figure 1A). This delay in age-dependent reproductive decline is also apparent in the increased capacity for aged *crh-1(-)* mutants to successfully produce progeny when mated past their reproductive prime (at day 7 of adulthood; Figure 1B).

**Figure 1.**
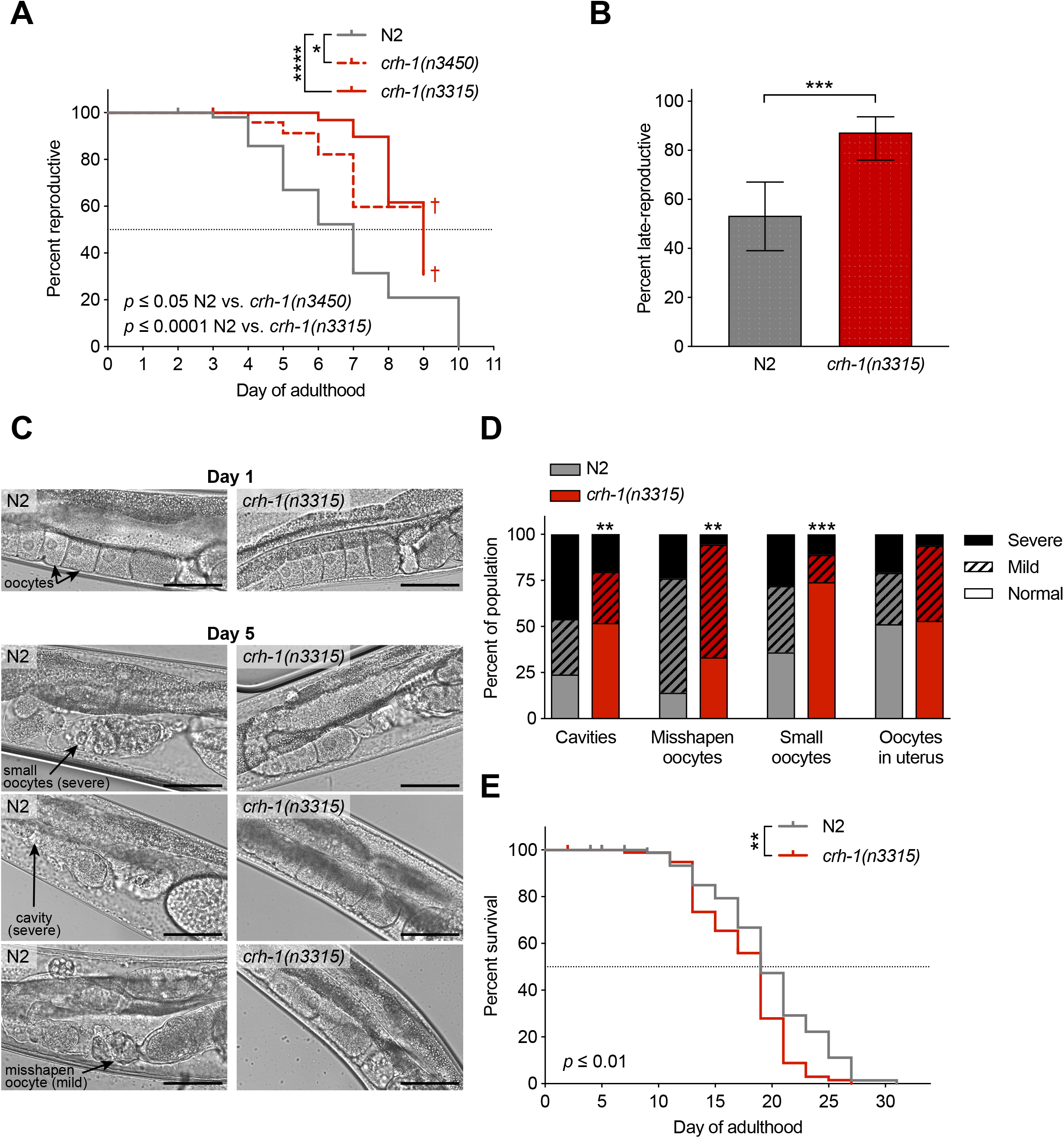
Loss of CREB signaling delays age-dependent reproductive decline. (A) Whole-body loss of CREB causes an extension of mated reproductive span compared to wild type (N2; n = 53), as seen in two *C. elegans* strains with different *crh-1* null or severe loss-of-function alleles (*crh-1(n3315)* and *crh-1(n3450)*, n = 25-33). † indicates a high matricide frequency. (B) A significantly higher percentage of the *crh-1(n3315)* population (n = 55) have the capacity to produce progeny when mated at day 7 of adulthood, compared to N2 (n = 45). Bars represent population means, error bars represent 95% confidence intervals. (C and D) Representative images (C) and scored oocyte morphology defects of worm populations (D) show significant improvements in oocyte quality for mated *crh-1(n3315)* worms (n = 54) on day 5 of adulthood, compared to N2 (n = 50). For reference, healthy oocytes from young (day 1 adult) worms are shown. Scale bars = 50 μm. (E) Loss of CREB does not extend lifespan. *crh-1(n3315)* worms (n = 96) have a slight reduction of lifespan, compared to N2 worms (n = 96). * *p* ≤ 0.05; ** *p* ≤ 0.01; *** *p* ≤ 0.001; **** *p* ≤ 0.0001.

Moreover, *crh-1(-)* mutation causes a significant improvement in the morphology of aging oocytes, a direct indicator of oocyte quality (Figure 1C). Young, healthy oocytes have a regular cuboidal shape, while by Day 5 of adulthood, wild-type oocytes have irregularly shaped or abnormally small oocytes, with prominent cavities between oocytes; by contrast, mated *crh-1(-)* oocytes on Day 5 appear significantly more youthful (Figures 1C, 1D). Therefore, CREB signaling regulates reproductive aging and oocyte quality maintenance.

Importantly, *crh-1(-)* mutants exhibit a significant delay in age-dependent reproductive decline without prolonged longevity. Consistent with previous results (Chen et al., 2016; Lakhina et al., 2015), *crh-1(-)* mutants are not long lived (Figure 1E); in fact, *crh-1(-)* mutants had a slightly reduced lifespan compared to wild type, as has been previously observed (Chen et al., 2016). Therefore, CREB is specifically involved in regulating oocyte quality maintenance without exerting corresponding effects on somatic tissue maintenance.

### Hypodermal CREB signaling regulates oocyte quality maintenance and reproductive aging

CREB’s ability to control diverse biological functions is likely enabled by transcriptional regulation of distinct target genes in different tissue types (Altarejos and Montminy, 2011)(Lakhina et al., 2015), therefore, we wished to determine in which tissue CREB acts to regulate reproductive aging. First, we evaluated the effects of rescuing *crh-1* expression in the neurons of *crh-1(-)* mutants, as neuronal CREB activity is critical for regulating long-term memory (Kauffman et al., 2010). However, expression of *crh-1* under a neuron-specific promoter that was demonstrated to be sufficient for CREB’s impact on long-term memory (Kauffman et al., 2010) and neuronal CRE-mediated transcription (Kimura et al., 2002; *i.e., crh-1(n3315);Pcmk-1∷crh-1β)* did not significantly change the rate of reproductive decline of *crh-1* mutants (Figure 2A).

**Figure 2.**
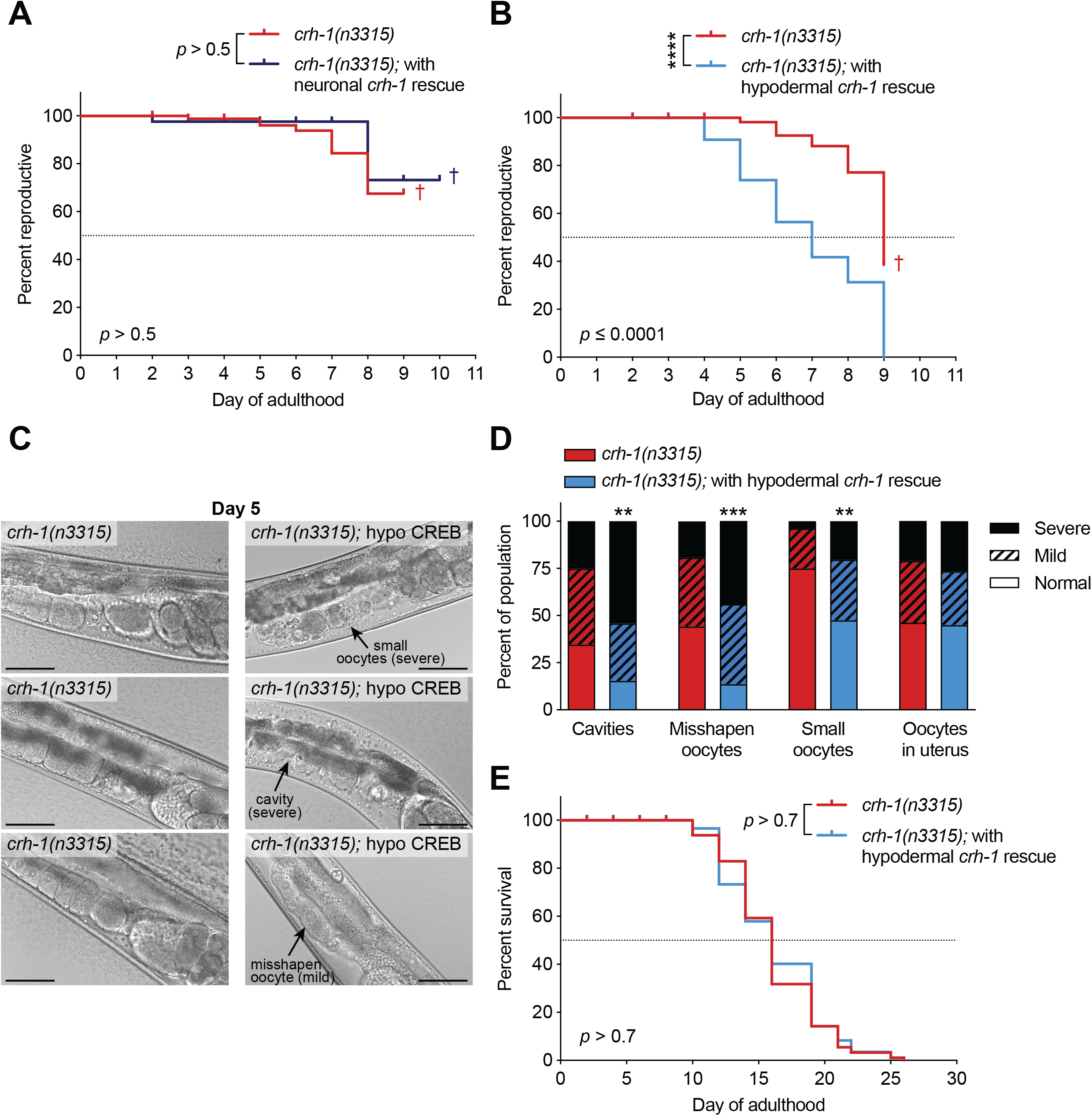
Hypodermal CREB signaling regulates oocyte quality maintenance and reproductive aging. (A) neuronal *crh-1* rescue *(crh-1(n3315);Pcmk-1∷crh-1β*; n = 41) does not significantly change the mated reproductive span of *crh-1(n3315)* worms (n = 93). (B) Rescuing *crh-1* expression in the hypodermis of *crh-1(-)* mutants reverses the reproductive span extension caused by whole-body CREB loss. hypodermal *crh-1* rescue *(crh-1(n3315);Pdpy-7∷crh-1β + PY37A1B.5∷crh-1β*, n = 83) significantly shortens the mated reproductive spans of long-reproducing *crh-1(n3315)* worms (n = 93). (C and D) Hypodermal CREB non cell-autonomously regulates oocyte quality maintenance. Representative images (C) and scored oocyte morphology defects of worm populations (D) show significantly worsened oocyte quality in mated *crh-1(n3315);Pdpy-7∷crh-1β* + *PY37A1B.5∷crh-1β* worms *(i.e., crh-1(n3315);* hypo CREB, n = 59) on day 5 of adulthood, compared to *crh-1(n3315)* worms (n = 52). Scale bars = 50 μm. (E) Hypodermal *crh-1* expression regulates reproductive aging independently of any effect on lifespan. *crh-1(n3315);Pdpy-7∷crh-1β* + *PY37A1B.5∷crh-1β* worms (n = 113) do not have a significantly different lifespan than *crh-1(n3315)* worms (n = 115). ** *p* ≤ 0.01; *** *p* ≤ 0.001; **** *p* ≤ 0.0001. † indicates a high matricide frequency.

We next considered the hypodermis, an epithelial tissue that we recently found to be important for metabolic functions such as nutrient storage (Kaletsky et al., 2018), in addition to its known roles regulating cuticle formation and function, molting, and body morphogenesis (Altun and Hall, 2009). Notably, the hypodermis is also the tissue in which TGF-β Sma/Mab signaling regulates growth (Wang et al., 2002) and reproductive aging (Luo et al., 2010). Gene Set Enrichment Analysis showed clear similarities between *crh-1* and TGF-β Sma/Mab loss-of-function mutants, as the transcriptional profile of *crh-1* worms (Lakhina et al., 2015) exhibits significant positive enrichment for genes altered in *sma-2* mutants (Luo et al., 2010; Figure S2A). In addition, *crh-1* mutants are small (Lakhina et al., 2015; Figure S1A), albeit not as small as TGF-β Sma/Mab loss-of-function mutants. Expressing *crh-1* under hypodermis-specific promoters *(i.e., crh-1(n3315);Pdpy-7∷crh-1β + PY37A1B.5∷crh-1β;* Kaletsky et al., 2018) increases their body length (Figure S2B). Therefore, the hypodermis seemed to be an excellent candidate tissue for mediating CREB’s regulation of reproductive aging.

We found that rescuing *crh-1* expression solely in the hypodermis was sufficient to rescue (shorten) *crh-1(-)* mutants’ extended mated reproductive span (Figure 2B). Furthermore, mated day 5 hypodermal *crh-1* rescue animals’ oocyte quality is significantly deteriorated compared to *crh-1(-)* mutants, as indicated by increasing frequencies of cavities between oocytes, irregularly shaped oocytes, and abnormally small oocytes (Figures 2C, 2D). Notably, hypodermal CREB signaling regulates oocyte quality and reproductive aging independently of any significant effects on lifespan (Figure 2E).

### Hypodermal CREB acts downstream of TGF-β Sma/Mab signaling to regulate the rate of reproductive decline

Considering that CREB and TGF-β Sma/Mab signaling in the hypodermis are both involved in the regulation of growth and reproductive aging, we wanted to determine the epistasis of these signaling pathways. As was previously shown, rescuing the expression of *sma-3* in the hypodermis of worms with a severe loss-of-function or null mutation of the R-SMAD *sma-3*, a TGF-β Sma/Mab receptor-regulated signaling transducer, is sufficient to rescue the very small body size of *sma-3(-)* mutants (Figure 3A; Wang et al., 2002). We found that rescuing *sma-3* expression in the hypodermis of *crh-1;sma-3* double mutants still lengthens body size, even when CREB is deleted (Figure 3A). This suggests two possibilities: either CREB acts upstream of hypodermal TGF-β Sma/Mab signaling to regulate growth, in which case directly rescuing hypodermal SMA-3 bypasses the requirement for CREB, or alternatively, these pathways regulate growth independently. Rescuing the expression of *sma-3* in the hypodermis also reverses the reproductive span extension of *sma-3* mutants (Figure 3B; Luo et al., 2010). However, rescuing hypodermal *sma-3* in *crh-1;sma-3* double mutants does not affect reproductive span (Figure 3C). This indicates that hypodermal TGF-β Sma/Mab signaling requires the presence of CREB to affect the rate of reproductive decline, either because these signaling pathways are working together, or because CREB acts downstream of TGF-β Sma/Mab signaling to regulate reproductive aging.

**Figure 3.**
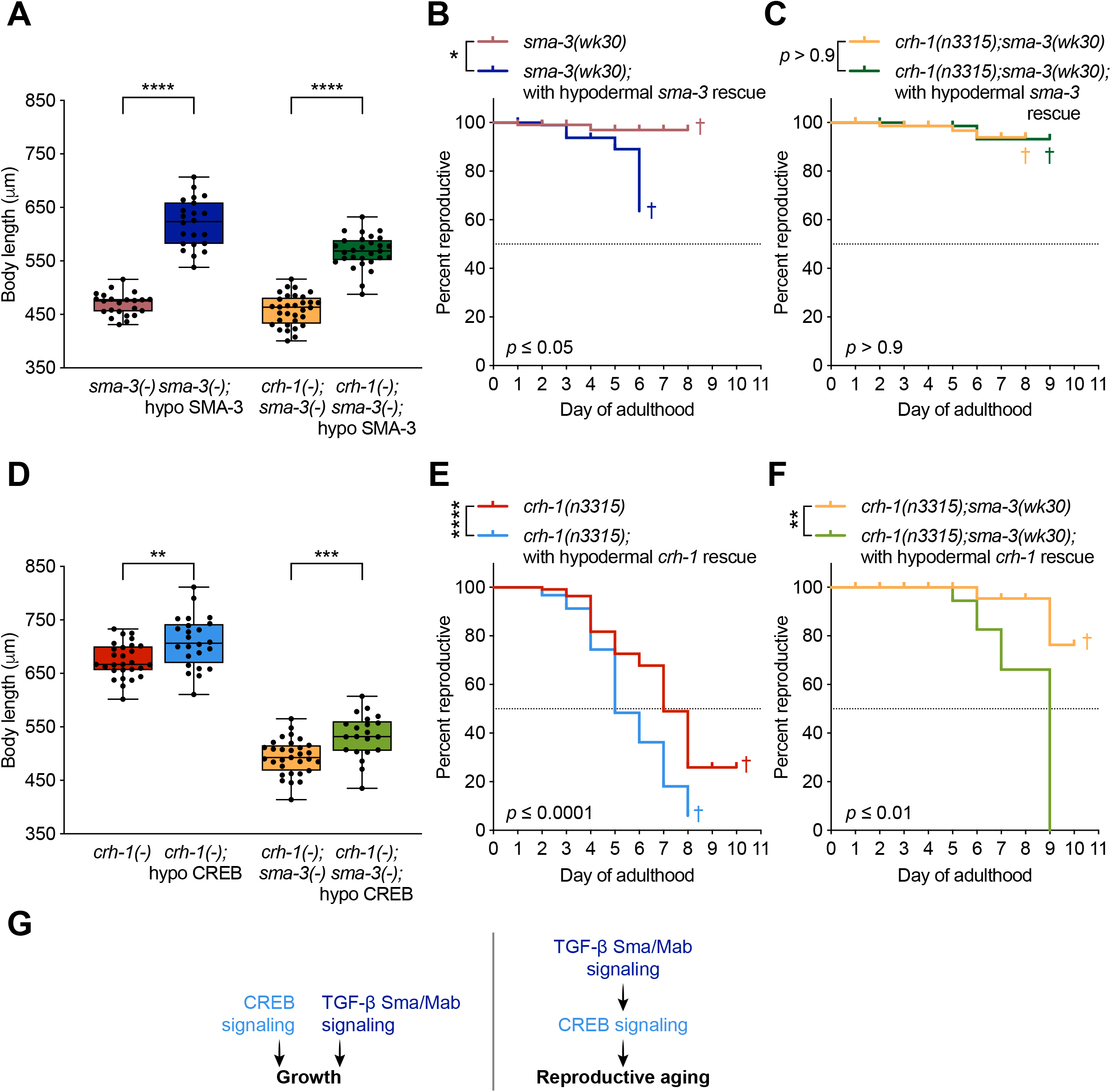
Hypodermal CREB acts downstream of TGF-**β** Sma/Mab signaling to regulate the rate of reproductive decline. (A) Hypodermal *sma-3* rescue is sufficient to increase growth of extremely small *sma-3(-)* mutants or double *crh-1(-);sma-3(-)* mutants. Hypodermal *sma-3* rescue *(sma-3(-);*hypo SMA-3, *i.e., sma-3(wk30);Pvha-7∷sma-3*, n = 22; and *crh-1(-);sma-3(-);*hypo SMA-3, *i.e., crh-1(n3315);sma-3(wk30);Pvha-7∷sma-3*, n = 29) significantly increases the body lengths of *sma-3(wk30)* worms (n = 23) and *crh-1(n3315);sma-3(wk30)* worms (n = 32), respectively. (B and C) Hypodermal *sma-3* rescue does not shorten the mated reproductive span of double *crh-1(-);sma-3(-)* mutants, indicating that hypodermal TGF-β Sma/Mab signaling requires CREB to regulate reproductive aging. Hypodermal *sma-3* rescue *(sma-3(wk30);Pvha-7∷sma-3*, n = 109) significantly shortens the mated reproductive span of long-reproducing *sma-3(wk30)* worms (n = 100; B), but *crh-1(n3315);sma-3(wk30)* mutants with hypodermal *sma-3* rescue *(crh-1(n3315);sma-3(wk30);Pvha-7∷sma-3*, n = 76) do not have significantly different mated reproductive spans than *crh-1(n3315);sma-3(wk30)* worms (n = 76; C). (D) Hypodermal *crh-1* rescue is sufficient to increase growth of small *crh-1(-)* mutants or extremely small double *crh-1(-);sma-3(-)* mutants. Hypodermal *crh-1* rescue *(crh-1(-);*hypo CREB, *i.e., crh-1(n3315);Pdpy-7∷crh-1β* + *PY37A1B.5∷crh-1β*, n = 24; and *crh-1(-);sma-3(-);*hypo CREB, *i.e., crh-1(n3315);sma-3(wk30);Pdpy-7∷crh-1β + PY37A1B.5∷crh-1β*, n = 21) significantly increases the body lengths of *crh-1(n3315)* worms (n = 28) and *crh-1(n3315);sma-3(wk30)* worms (n = 31), respectively. (E and F) Hpodermal *crh-1* rescue shortens the mated reproductive spans of both *crh-1(-)* mutants and double *crh-1(-);sma-3(-)* mutants, indicating that CREB does not require hypodermal TGF-β Sma/Mab signaling to regulate reproductive aging. Hypodermal *crh-1* rescue *(crh-1(-);*hypo CREB, *i.e., crh-1(n3315);Pdpy-7∷crh-1β* + *PY37A1B.5∷crh-1β*, n = 92; and *crh-1(-);sma-3(-);*hypo CREB, *i.e., crh-1(n3315);sma-3(wk30);Pdpy-7∷crh-1β* + *PY37A1B.5∷crh-1β*, n = 105) significantly shortens the mated reproductive spans of *crh-1(n3315)* worms (n = 111) and *crh-1(n3315);sma-3(wk30)* worms (n = 126), respectively. (G) Summarized interpretation of collective results from A-F. Boxes represent 25th to 75th percentiles, with lines at the means, points represent individual worms, whiskers represent minimum and maximum values. * *p* ≤ 0.05; ** *p* ≤ 0.01; *** *p* ≤ 0.001; **** *p* ≤ 0.0001. † indicates a high matricide frequency.

To distinguish between these possibilities, we performed the converse set of epistasis experiments. Expressing *crh-1* in the hypodermis lengthens the body sizes of both *crh-1* mutants and *crh-1;sma-3* double mutants (Figure 3D), which demonstrates that CREB and TGF-β Sma/Mab signaling independently regulate body growth. Moreover, similar to hypodermal *crh-1* rescue in *crh-1* mutants (Figures 2B, 3E), rescuing *crh-1* expression in the hypodermis of *crh-1;sma-3* double mutants is sufficient to shorten their extended mated reproductive span (Figure 3F). This suggests that CREB activity in the hypodermis may be regulated by TGF-β Sma/Mab signaling, and that the ensuing change in hypodermal CREB activity is responsible for regulating reproductive aging (Figure 3G).

### CREB’s hypodermal transcriptional targets include a Hedgehog-related signaling factor that affects reproductive aging

Our next goal was to delineate CREB’s biological functions in the hypodermis by identifying and characterizing its hypodermis-specific transcriptional target gene set. We used our chilled chemo-mechanical disruption technique followed by fluorescence-activated cell sorting (Kaletsky et al., 2016) to isolate and purify GFP-labeled hypodermal tissue samples from *crh-1a* nd wild-type *C. elegans* on day 1 of adulthood *(i.e., crh-1(n3315);Pdpy-7∷gfp* and *N2;Pdpy-7∷gfp)*, and performed RNA sequencing of the purified samples. Compared to wild type, 224 genes were significantly downregulated in the hypodermis of *crh-1* mutants (Figures 4A, 4B, Table S1; adjusted p-value of 0.05), and 101 genes were significantly upregulated in *crh-1* hypodermal tissue (Figures 4A, 4C, Table S1). The majority of these significantly altered genes had been previously determined to be expressed in hypodermal tissue (Figure 4D; Kaletsky et al., 2018). Strikingly, many of the genes downregulated in the hypodermis of *crh-1* animals are associated with metabolism (Figure 4B, Table S1), and gene ontology (GO) analysis of this gene set highlights the enrichment of metabolic processes such as carboxylic acid metabolism, small molecule metabolism, oxidation-reduction activity, cofactor metabolic processes, as well as amino acid, fatty acid, and glucose metabolism (Figure 4E, Table S2), suggesting that CREB plays an integral role in regulating metabolic functions and energy homeostasis in *C. elegans* hypodermal tissue. By contrast, many of the genes upregulated in the hypodermis of *crh-1* mutants are associated with GO terms related to molting and cuticle structure and development (Figure 4F, Table S2). Collectively, the hypodermis-specific transcriptional targets of CREB reveal that this transcription factor is involved in governing many key biological functions of the *C. elegans* hypodermis, including development and metabolism.

**Figure 4.**
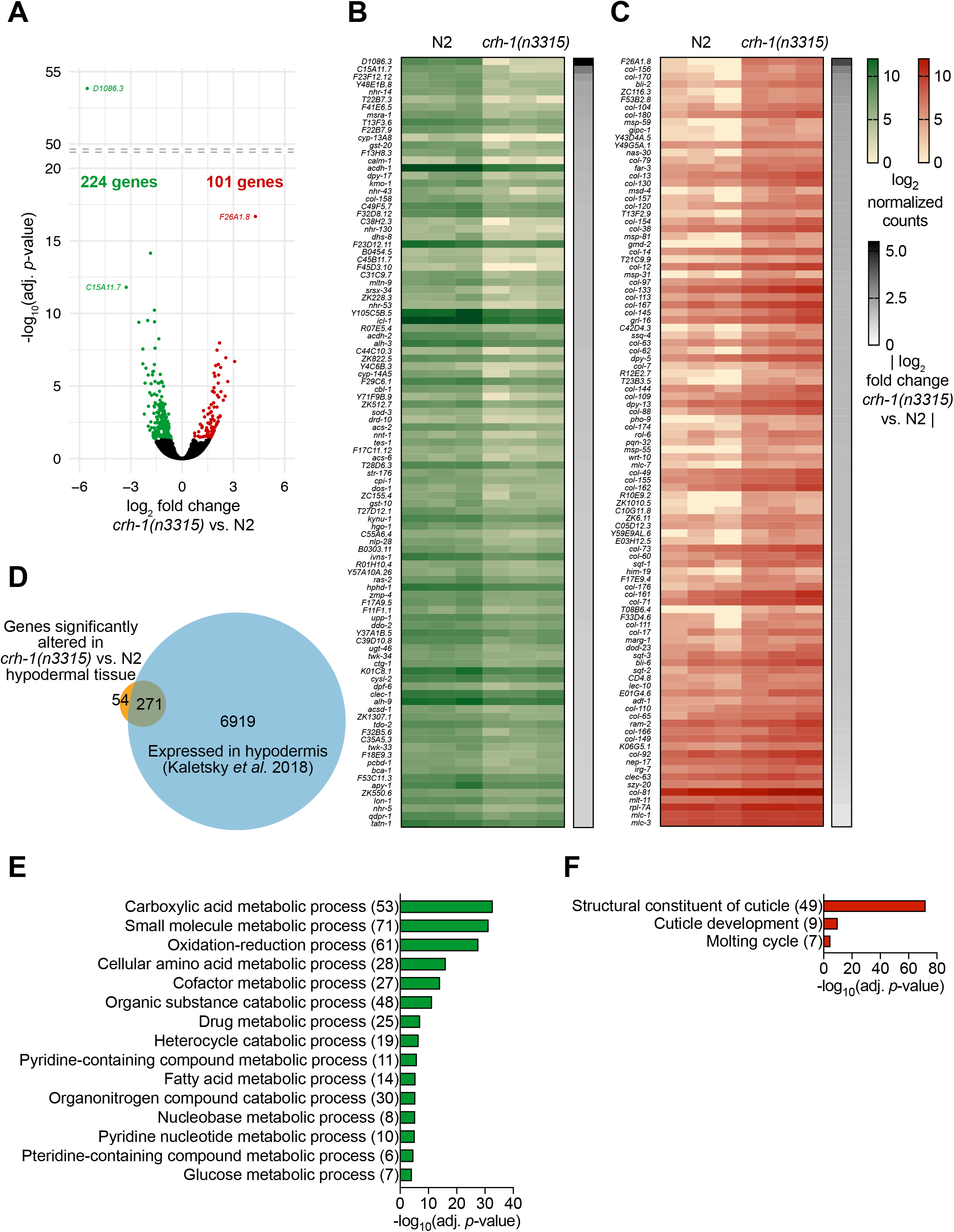
Identification of CREB’s hypodermis-specific transcriptional targets. (A) Volcano plot showing 224 genes that were significantly downregulated (green) and 101 genes that were significantly upregulated (red) in hypodermal tissue from *crh-1(n3315);Pdpy-7∷gfp* worms, compared to *N2;Pdpy-7∷gfp.* Colored points indicate genes that are significantly altered (adj. or adjusted p-value ≤ 0.05) from three independent collections of day 1 adult worms. (B and C) Top 101 significantly downregulated (B) and upregulated (C) genes, ranked by |fold change|, of three independent expression comparisons of hypodermal tissue from day 1 *crh-1(n3315);Pdpy-7∷gfp* worms versus *N2;Pdpy-7∷gfp.* For complete lists of significantly altered genes, see Table S1. (D) More than 80% of the 325 genes significantly altered in hypodermal tissue from *crh-1(n3315);Pdpy-7∷gfp* versus *N2;Pdpy-7∷gfp*, were previously identified as being expressed in *C. elegans* hypodermal tissue (Kaletsky et al., 2018). (E and F) Top biological processes associated with the genes downregulated in the hypodermis of *crh-1(-)* worms (E), and top biological processes and molecular functions associated with the genes upregulated in the hypodermis of *crh-1(-)* worms (F). Numbers in parentheses signify the number of genes contributing to enrichment for each GO term. Adj. *p*-value ≤ 0.0001 for top GO terms, and overlapping GO terms were excluded where necessary *(i.e.* child and parent terms were not both included). For complete lists of significant GO terms, see Table S2.

To determine how CREB activity in the hypodermis regulates reproductive aging cell non-autonomously, we examined the effect of hypodermal CREB targets on late-mating capacity. We tested 14 candidate genes that were significantly down- or up-regulated in the hypodermis of day 1 adult *crh-1* mutants, including candidates that were highly ranked based on fold-change (Figures 4B, 4C), significantly altered in the whole bodies of *sma-2* TGF-β Sma/Mab signaling mutant L4 larvae (Luo et al., 2010; Figure S3), and/or whose predicted functions involve intercellular signaling (Table S1). Compared to wild type, *crh-1* mutants have an increased capacity for progeny production when mated on day 7 of adulthood (Figures 1B, 5A-D); if a *crh-1*-upregulated gene is required, then knocking down that gene in *crh-1(-)* mutants will worsen their late-mating success. RNA interference (RNAi)-mediated knockdown of most candidate genes did not alter *crh-1* mutants’ late-mating capability (Figures 5A, 5B), and knockdown of candidate *crh-1*-downregulated genes did not significantly improve the late-mating capacity of wild type animals (Figures 5C, 5D). However, knockdown of *wrt-10*, which encodes a Hedgehog-related signaling factor, significantly impaired aged *crh-1* mutants’ ability to produce progeny (Figure 5A).

**Figure 5.**
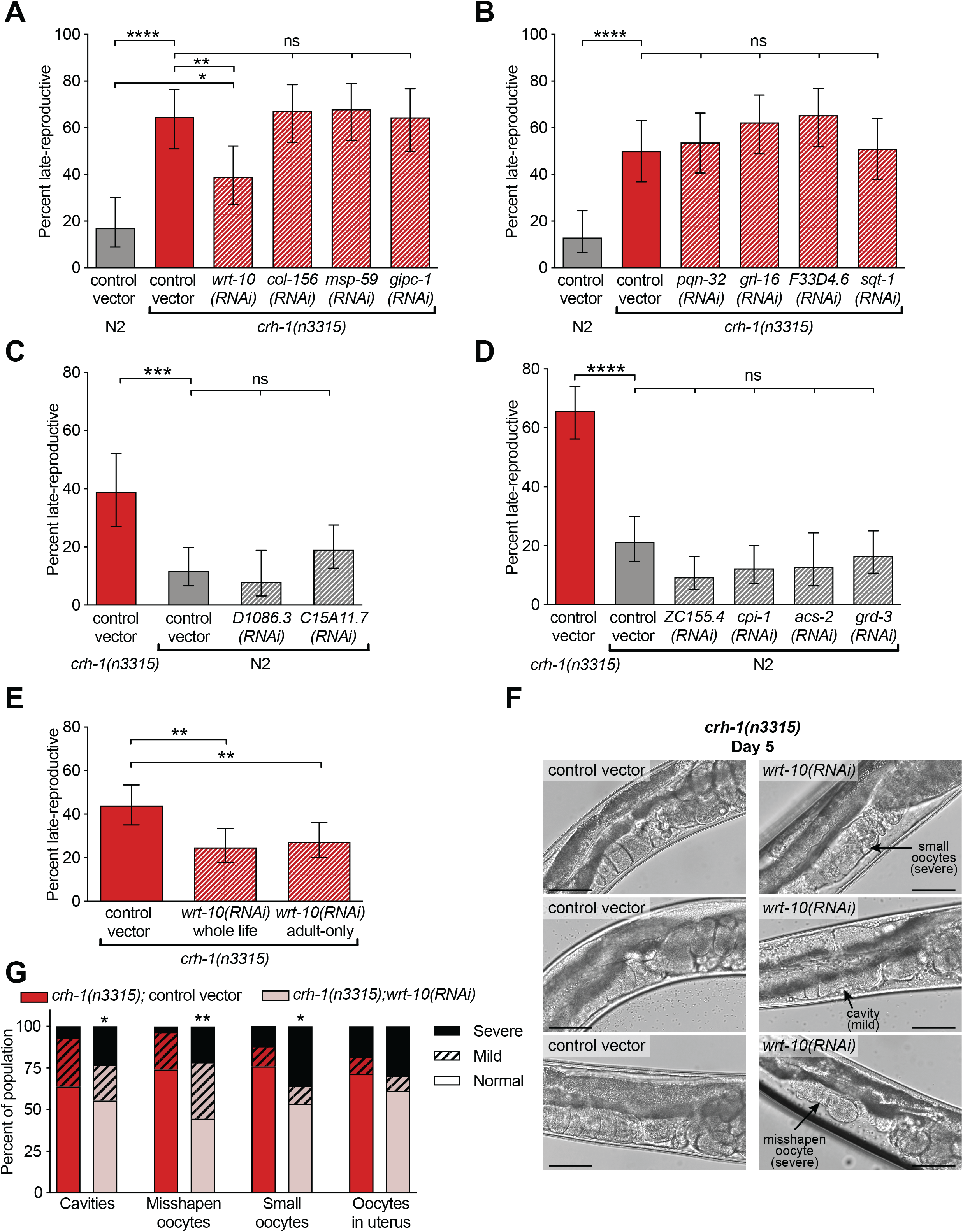
Characterization of CREB’s hypodermal transcriptional targets reveals that a Hedgehog-related signaling factor regulates reproductive aging. (A and B) Upregulation of hedgehog-related *wrt-10* expression contributes to the superior late-mating capacity of *crh-1(-)* worms. Whole-life knockdown of most of the tested *crh-1(-)* upregulated genes did not significantly worsen the late-mating success of *crh-1(n3315)* worms, with the exception of *wrt-10* (A). (C and D) Whole-life knockdown of the tested *crh-1(-)* downregulated genes did not significantly improve the late-mating success of N2 worms. (E) Exposure to *wrt-10* RNAi from day 1 of adulthood onward (“adult-only”) is sufficient to worsen the late-mating capacity of *crh-1(n3315)* worms mated at day 7 of adulthood, to a similar degree as *crh-1(n3315)* worms exposed to *wrt-10* RNAi from egg stage onward (“whole life”). In A-E, each panel is a separate experiment, bars represent population means, and error bars represent 95% confidence intervals. n = 45-117. Bonferroni corrections were applied to adjust for multiple comparisons (ns = nonsignificant, *i.e.*, greater than Bonferroni-adjusted α-value). * *p* ≤ 0.025; ** *p* ≤ 0.01; *** *p* ≤ 0.001; **** *p* ≤ 0.0001; (A,B,D) ns *p* > 0.01; (C) ns *p* > 0.017. (F and G) WRT-10 regulates oocyte quality maintenance. Representative images (F) and scored oocyte morphology defects of worm populations (G) show significantly worsened day 5 oocyte quality in mated *crh-1(n3315)* with whole-life *wrt-10* knockdown (n = 56), compared to *crh-1(n3315)* worms exposed to control vector (n = 58). * p ≤ 0.05; ** *p* ≤ 0.01.

WRT-10 is a member of the large family of Hedgehog-related signaling factors in *C. elegans* (Aspöck et al., 1999). Its contribution to regulation of reproductive aging may be distinct, as knockdown of *grd-3* or *grl-16*, which encode two other Hedgehog-related signaling factors with altered expression levels in *crh-1* hypodermal tissue, did not significantly change late-mating success (Figures 5B, 5D). While the roles of *C. elegans* Hedgehog-related proteins have not be fully characterized, many contribute to such key developmental functions as molting, growth, and morphogenesis (Zugasti et al., 2005). Importantly, we found that adult-only reduction of *wrt-10* reduces the capacity of *crh-1* mutants to successfully produce progeny when mated at day 7 of adulthood (Figure 5E), indicating that WRT-10 acts post-developmentally to regulate reproductive aging. Furthermore, knockdown of *wrt-10* worsens the oocyte quality of day 5 mated *crh-1* mutants, as evidenced by higher incidences of inter-oocyte cavities, irregularly shaped oocytes, or abnormally small oocytes (Figures 5F, 5G). Elevated *wrt-10* is required for the enhanced oocyte quality maintenance and delayed reproductive aging of *crh-1* mutants.

### Hypodermal overexpression of Hedgehog-related *wrt-10* delays age-dependent reproductive decline and oocyte quality deterioration

To ascertain whether hypodermis-derived WRT-10 is critical for regulating the rate of reproductive decline, we tested whether overexpression of *wrt-10* specifically in the hypodermis could slow reproductive aging in a wild-type background. Previous promoter∷GFP fusion experiments showed that *wrt-10* is expressed in the hypodermis from the embryo stage through adulthood (Hao et al., 2006). We found that increasing hypodermal *wrt-10* expression under hypodermis-specific promoters *(Pdpy-7∷wrt-10 + PY37A1B.5∷wrt-10)* is sufficient to significantly extend the mated reproductive span of wild-type animals (Figure 6A). Moreover, day 5 mated worms with hypodermal *wrt-10* overexpression exhibit clear improvements in oocyte quality compared to wild type worms, as indicated by the reduced frequencies of cavities between oocytes, irregularly shaped oocytes, or abnormally small oocytes (Figures 6B, 6C). Therefore, hypodermis-expressed *wrt-10* plays a role in cell non-autonomous regulation of oocyte quality and reproductive aging.

**Figure 6.**
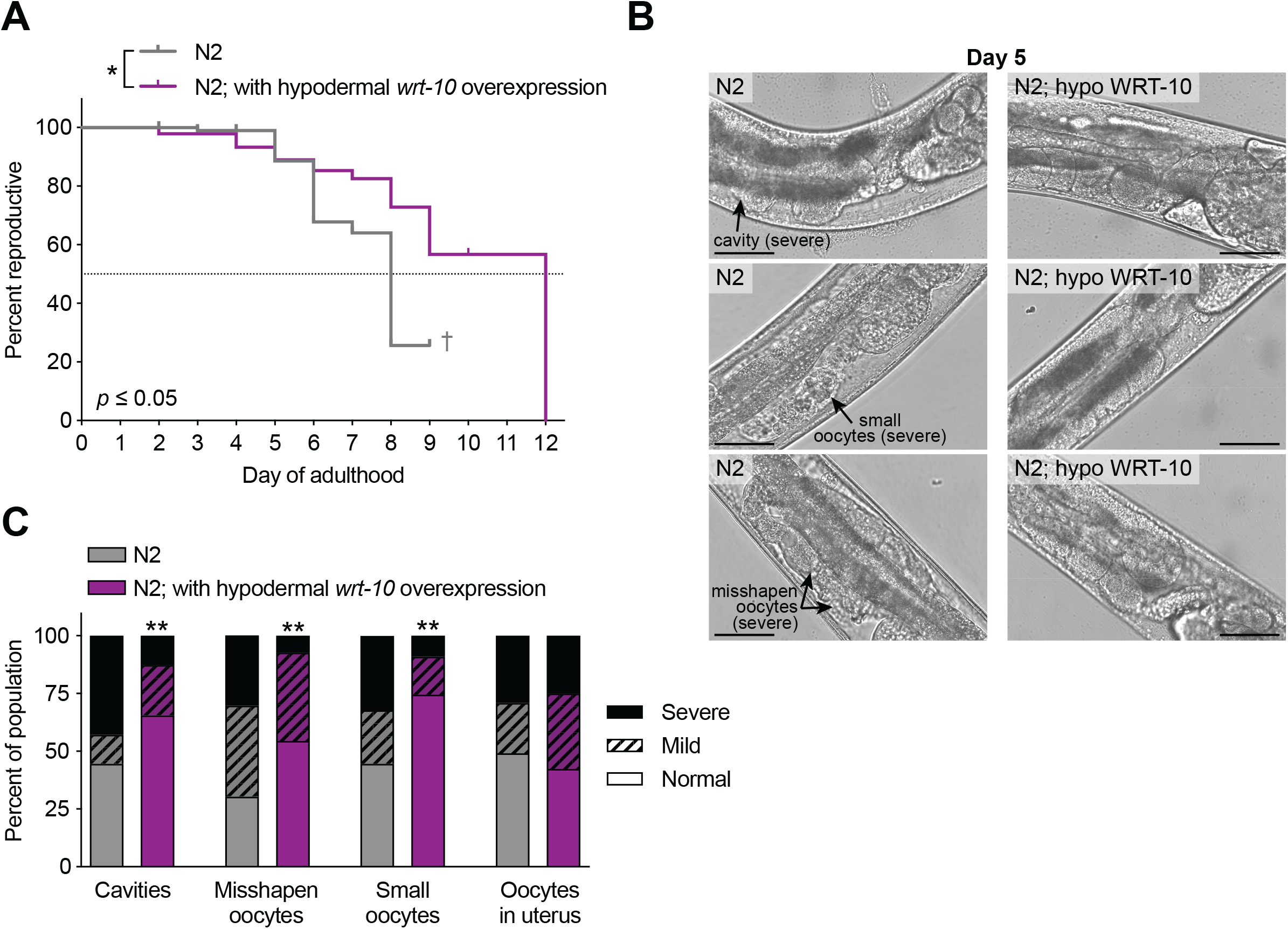
Hypodermal overexpression of Hedgehog-related *wrt-10* delays age-dependent reproductive decline and oocyte quality deterioration. (A) Overexpressing *wrt-10* specifically in the hypodermis of wild-type worms delays reproductive decline. Worms with hypodermal overexpression of *wrt-10 (N2;Pdpy-7∷wrt-10 + PY37A1B.5∷wrt-10*, n = 93) have significantly longer mated reproductive spans than wild type (n = 98). † indicates a high matricide frequency. (B and C) Hypodermal WRT-10 non cell-autonomously regulates oocyte quality maintenance. Representative images (B) and scored oocyte morphology defects of worm populations (C) show significantly improved oocyte quality in mated worms with hypodermal overexpression of *wrt-10 (N2;Pdpy-7∷wrt-10 + PY37A1B.5∷wrt-10)* on day 5 of adulthood, compared to wild type. n = 55-56. Scale bars = 50 μm.

### Patched-related receptors in the germ cells regulate reproductive aging and oocyte quality maintenance

WRT-10 is likely to be a secreted molecule, based on its predicted N-terminal protein export sequence (Aspöck et al., 1999). Therefore, we hypothesized that WRT-10 acts as a signaling factor communicating between the hypodermis and the reproductive system. The Patched receptor is the core receptor for the Hedgehog ligand in *Drosophila melanogaster* and vertebrates, so it seemed possible that *C. elegans* Patched-related receptors in the germline could mediate the Hedgehog-related WRT-10 signal. In fact, the Patched receptor PTC-1 has a crucial role in the *C. elegans* reproductive system. PTC-1 protein and *ptc-1* gene expression is localized to the germline and developing oocytes, and deletion or whole-body RNAi-mediated knockdown of *ptc-1* causes a high incidence of sterility and multinucleate oocytes, suggesting an involvement in regulating germline cytokinesis (Kuwabara et al., 2000). Whole-body RNAi-mediated knockdown of the Patched-related receptor *ptr-2* also causes embryonic arrest (Zugasti et al., 2005).

Therefore, we tested how germline-specific RNAi-mediated knockdown of four of the Patched-related receptors that have especially elevated expression in oocytes (Stoeckius et al., 2014) and oogenic gonads (Ortiz et al., 2014) affects reproductive aging of worms with hypodermal *wrt-10* overexpression *(i.e., rde-1(mkc36);Psun-1∷rde-1;Pdpy-7∷wrt-10 + PY37A1B.5∷wrt-10*, see (Zou et al., 2019)). We used adult-only RNAi exposure to avoid developmental effects, and mated and maintained hermaphrodites on control vector from day 5 onward to preclude potential effects of RNAi on males. *rde-1(mkc36);Psun-1∷rde-1* worms on either RNAi or control vector had a notably short period of adulthood during which they were capable of progeny production when mated, so we performed these late-mating assays on day 5 of adulthood. We found that germline-specific knockdown of *ptc-1* or *ptr-2* caused a pronounced reduction in late-mating capacity, compared to worms exposed to control vector (Figure 7A).

**Figure 7.**
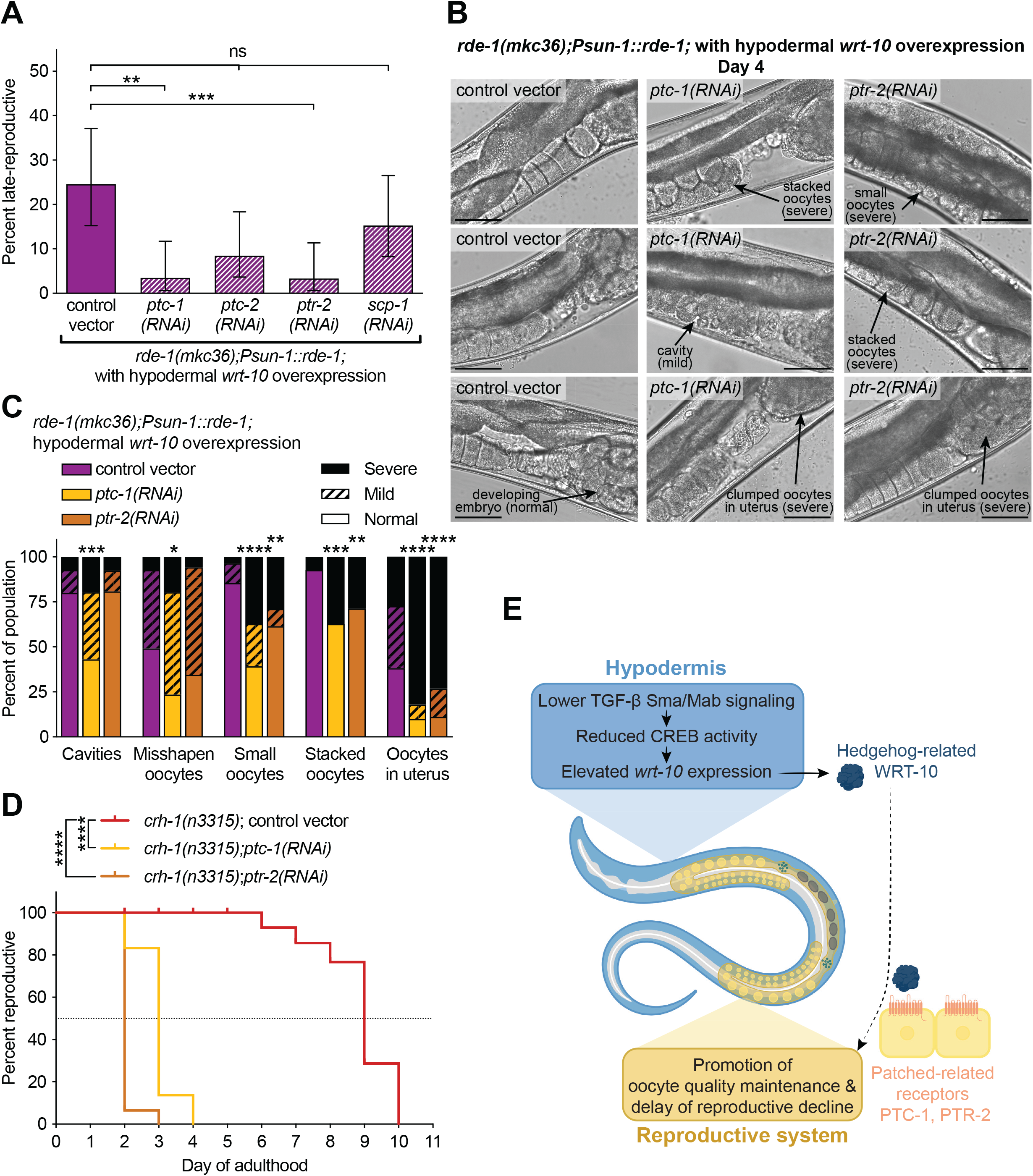
Patched-related receptors in the germ cells regulate reproductive aging and oocyte quality maintenance. (A) Adult-only, germline-specific knockdown of Patched-related receptors *ptc-1* and *ptr-2* worsens the late-mating success of worms overexpressing hypodermal *wrt-10.* In hypodermal *wrt-10* overexpression worms that are specifically sensitive to RNAi in the germline (*rde-1(mkc36);Psun-1∷rde-1;Pdpy-7∷wrt-10 + PY37A1B.5∷wrt-10*, see (Zou et al., 2019)), adult-only (day 1 of adulthood onward) exposure to *ptc-1* or *ptr-2* RNAi caused a significant reduction in the capacity of worms to produce progeny when mated at day 5 of adulthood; exposure to *ptc-2* or *scp-1* RNAi did not have a pronounced effect on late-mating capacity. Bars represent population means, error bars represent 95% confidence intervals. n = 57-60. (B and C) Patched-related receptors *ptc-1* and *ptr-2* act in the germline to regulate oocyte quality maintenance. Representative images (B) and scored oocyte morphology defects of populations of day 4 hypodermal *wrt-10* overexpression worms that are RNAi-sensitive in the germline (*rde-1(mkc36);Psun-1∷rde-1;Pdpy-7∷wrt-10* + *PY37A1B.5∷wrt-10)* (C) show significantly worsened oocyte quality in worms exposed to *ptc-1* or *ptr-2* RNAi from day 1 of adulthood onward. n = 51-55. Scale bars = 50 μm. (D) Adult-only (day 1 of adulthood onward) knockdown of Patched-related receptors *ptc-1* (n = 84) and *ptr-2* (n = 92) significantly shortens the mated reproductive span of long-reproducing *crh-1(n3315)* worms (n = 76). (E) Model of proposed signaling axis wherein the transcription factor CREB acts downstream of TGF-β Sma/Mab signaling in the hypodermis to control reproductive aging. CREB exerts these reproductive effects at least in part by regulating the hypodermal mRNA levels of a Hedgehog-related signaling factor, *wrt-10.* Elevated levels of hypodermis-derived WRT-10 improves oocyte quality maintenance and extends reproductive span, potentially (dotted line) via the Patched-related receptors PTC-1 and/or PTR-2 in the germline. Bonferroni corrections applied to adjust for multiple comparisons (ns = nonsignificant, *i.e.*, greater than Bonferroni-adjusted α-value); * *p* ≤ 0.025; ** *p* ≤ 0.01; *** *p* ≤ 0.001; **** *p* ≤ 0.0001; ns *p* > 0.0125 in comparisons to control vector.

Furthermore, adult-only, germline-specific knockdown of either *ptc-1* or *ptr-2* significantly worsens the oocyte quality of day 4 mated hypodermal *wrt-10* overexpression worms. We observed a very high occurrence of clumped unfertilized oocytes (and likely other cellular material) in the uteruses of *ptc-1(RNAi)* and *ptr-2(RNAi)* animals (Figures 7B, 7C), suggesting an accumulation of defective oocytes and/or arrested embryos consistent with previously observations of multinucleate oocytes or embryonic arrest (Kuwabara et al., 2000; Zugasti et al., 2005), or other oocyte deficiencies. In addition, *ptc-1(RNAi)* worms have severe phenotypes associated with age-dependent oocyte quality deterioration, namely, higher incidences of inter-oocyte cavities, irregularly shaped oocytes, and abnormally small oocytes (Figures 7B, 7C). *ptr-2(RNAi)* worms displayed an increased frequency of abnormally small oocytes, and both *ptc-1(RNAi)* and *ptr-2(RNAi)* showed disarray in the developing oocytes of the proximal gonad, as evidenced by “stacking” of oocytes in multiple fields of view (Figures 7B, 7C). These results suggest that the Patched-related receptors PTC-1 and PTR-2 have cell-autonomous roles in regulating oocyte quality maintenance with age. In addition, knockdown of either *ptc-1* or *ptr-2* dramatically reduces the mated reproductive span of *crh-1* mutants (Figure 7D), establishing their critical functions in the reproductive system of long-reproducing *crh-1* mutant worms. Therefore, these receptors may represent a germline-specific signaling hub that allows for hypodermis-derived WRT-10 to regulate the rate of age-related reproductive decline (Figure 7E).

## Discussion

Given the high energetic demands of reproduction, it is essential for organisms to be able to adjust reproductive function in response to nutrient levels. For instance, under conditions of nutrient depletion, it is advantageous to reduce immediate progeny production in favor of extending cell maintenance and delaying reproductive decline (Templeman and Murphy, 2018). In this study, we revealed an intra- and inter-tissue signaling axis by which the relationship between energy balance and reproductive function can be coordinated. Our previous (Luo et al., 2010) and current findings highlight the role of the *C. elegans* hypodermis in determining the rate of reproductive decline; this tissue is clearly important in *C. elegans* for coordinating major biological functions relevant to energy homeostasis. Here, we show that: 1) hypodermal TGF-β Sma/Mab signaling requires the CREB transcription factor to regulate reproductive aging; 2) hypodermal CREB activity controls oocyte quality maintenance and reproductive aging, at least in part via the Hedgehog-related gene *wrt-10;* and 3) levels of hypodermis-derived WRT-10 signaling factor determine the rate of reproductive decline and the maintenance of oocyte quality with age, potentially by interacting with Patched-related receptors in the germline and oocytes.

### TGF-β Sma/Mab and CREB signal in the Hypodermis to separately regulate growth and reproduction

TGF-β Sma/Mab signaling, a *C. elegans* growth factor pathway homologous to the mammalian BMP pathway (Savage-Dunn and Padgett, 2017), plays an integral role in promoting energy-expensive functions such as lipid storage (Clark et al., 2018; Yu et al., 2017), body growth (Savage-Dunn et al., 2003), and progeny production (Luo et al., 2010). All of these functions are regulated by the actions of the TGF-β Sma/Mab pathway in the hypodermis (Clark et al., 2018; Luo et al., 2010; Wang et al., 2002), but they can be dissociated from each other (Luo et al., 2010; Yu et al., 2017). Our results point to one downstream mechanism by which hypodermal TGF-β Sma/Mab signaling separately controls these biological processes: while growth is discretely regulated by CREB and the TGF-β Sma/Mab pathway, hypodermal TGF-β Sma/Mab signaling requires the presence of CREB to alter the rate of reproductive decline, indicating that CREB enables hypodermal TGF-β Sma/Mab signaling to cell non-autonomously control reproductive aging. In vertebrates, CREB phosphorylation and transcriptional activity is promoted by numerous growth factor signals (Mayr and Montminy, 2001); TGF-β and BMP ligands can stimulate CREB phosphorylation (Gupta et al., 1999; Kramer et al., 1991; Lee and Chuong, 1997; Zhang et al., 2004), and CREB is required for a subset of the signaling events and responses that are mediated by Smad proteins involved in BMP signal transduction (Ionescu et al., 2004; Ohta et al., 2008). Thus, one of the effects of hypodermal TGF-β Sma/Mab signaling could be to induce CREB activity to promote reproduction.

While CREB is a highly conserved transcription factor that directs many diverse functions, its role in regulating age-related oocyte quality and reproductive decline has not been previously described in any organism. It is possible that CREB has the capacity to independently control diverse functions by regulating distinct target gene sets, depending on the tissue in which it acts and the other signaling systems that are involved. Upregulated genes in the hypodermal tissue of long-reproducing *crh-1* mutants included Hedgehog-related WRT-10, a signaling factor which is likely involved in relaying information from the hypodermis—a central metabolic tissue—to the reproductive system (Figure 7E). Importantly, hypodermis-specific overexpression of *wrt-10* is sufficient to promote oocyte quality maintenance and slow reproductive decline. Thus, decreased TGF-β Sma/Mab signaling ultimately delays age-dependent reproductive decline through CREB’s decreased activity in the hypodermis and subsequent WRT-10 signaling to oocytes (Figure 7E).

### Hedgehog-related/Patched-related signaling

The Hedgehog signaling system is a dynamic, complex network that encompasses more than the canonical signaling pathway, and has regulatory roles beyond its well-known functions in controlling pattern formation and cell proliferation (Robbins et al., 2012). For instance, in *Drosophila* larvae, a circulating lipoprotein-associated form of the Hedgehog protein is involved in coordinating metabolic processes, growth, and other physiological changes in response to nutrient deprivation, acting in part through a noncanonical pathway (Rodenfels et al., 2014). In *C. elegans*, there is an expansive group of Hedgehog-related families that are believed to share a common ancestor with the Hedgehog molecules of other phyla (Aspöck et al., 1999). While many of these Hedgehog-related proteins—including WRT-10—are predicted to be secreted signaling factors (Aspöck et al., 1999), neither their biological functions nor the signaling cascades by which they exert their effects have been fully characterized. *C. elegans* do have a superfamily of receptors that are homologous or related to the Patched receptor (the core Hedgehog receptor in *Drosophila* and vertebrates), but they do not appear to have homologs to some of the other components in the canonical signaling pathway (Kuwabara et al., 2000). Shared biological functions indicate that *C. elegans* Hedgehog-related proteins and Patched-related receptors may function cooperatively in the same network, perhaps through a ligand/receptor-activated signaling cascade, or via a Patched-related receptor response to lipids transported by Hedgehog-related factors (Kuwabara et al., 2000; Zugasti et al., 2005).

Our findings suggest that production of the Hedgehog-related signaling factor WRT-10 in hypodermal tissue leads to reproductive aging effects that could be mediated, at least in part, by Patched-related receptors in the germline and/or oocytes. *wrt-10* has been recently shown to contribute to *glp-1/*Notch’s regulation of germline stem cell maintenance (Roy et al., 2018), supporting the concept that this Hedgehog-related protein has key functions in the reproductive system. Several Patched-related receptors are highly expressed in oocytes and oogenic gonads (Stoeckius et al., 2014), and we found that germline-specific knockdown of *ptc-1* or *ptr-2* significantly impairs the late-mating capacity and oocyte quality of worms with hypodermal *wrt-10* overexpression (Figure 7). *ptc-1* or *ptr-2* knockdown have very severe impacts on reproductive phenotypes, which indicates that their functions likely extend beyond mediating WRT-10 signaling (unsurprising considering their critical roles in germline cytokinesis (Kuwabara et al., 2000) and embryonic development (Zugasti et al., 2005)). However, it is clear that the hypodermis-derived Hedgehog-related WRT-10 signaling factor and germline-specific Patched-related receptors are both involved in regulating reproductive function and oocyte quality.

In summary, here we have delineated a hypodermis-to-oocyte signaling axis which connects three major signaling pathways—the TGF-β Sma/Mab pathway, CREB-dependent transcription regulation, and Hedgehog-related/Patched-related signaling—in the regulation of oocyte quality maintenance and age-related reproductive decline. We show that CREB lies at a central signaling junction expressly responsible for directing the rate of reproductive aging, without exerting coordinated effects on somatic tissue aging or lifespan. This TGF-β/CREB/Hh axis integrates signaling across tissues to coordinate metabolic function with reproductive output and quality control with age.

## Acknowledgments

We thank Jasmine Ashraf and Daniel Xu for their contributions towards experiments, Christina DeCoste and Katherine Rittenbach in the Princeton Molecular Biology Flow Cytometry Resource Facility (partially supported by the Cancer Institute of New Jersey Cancer Center Support Grant; P30CA072720), the Princeton Genomics Core Facility, Patricia Kuwabara for providing the *ptc-1* RNAi clone, the *Caenorhabditis* Genetics Center (CGC) for strains, and members of the Murphy Lab for comments on the manuscript. N.M.T. was supported by a Canadian Institutes of Health Research Banting Postdoctoral Fellowship. The study was supported by the HHMI Faculty Scholar Program (AWD1005048), the Glenn Foundation for Medical Research (GMFR CNV1001899), as well as NIH New Innovator (1DP20D004402-01), DP1 Pioneer (NIGMS 1DP20D004402-01) and March of Dimes awards to C.T.M.

## Author Contributions

W.K. made the initial observations in preliminary experiments; N.M.T. and C.T.M. were involved in experimental design; N.M.T., V.C., and R.K. performed experiments and analyzed data; N.M.T. and C.T.M. wrote and edited the manuscript.

## STAR Methods

### CONTACT FOR REAGENT AND RESOURCE SHARING

Further information and requests for resources and reagents should be directed to and will be fulfilled by the Lead Contact, Coleen Murphy.

### EXPERIMENTAL MODEL AND SUBJECT DETAILS

#### *C. elegans* genetics and maintenance

The following strains were used in this study: the N2 Bristol strain as wild-type worms, *fog-2(q71) V* males for mating experiments, *crh-1(n3315), crh-1(n3450), crh-1(n3315);Pcmk-1∷crh-1β, crh-1(n3315);wqEx56[Pdpy-7∷crh-1β∷unc-54 3’ UTR* + *pY37A1B.5∷crh-1β∷unc-54 3’UTR + Pmyo-2∷mCherry]* (with *crh-1(n3315)* worms lacking the extrachromosomal array, from the same population, as the experimental comparison group), *sma-3(wk30) III;wqExl[Pvha-7∷sma-3 + Pmyo-2∷mCherry]* (with *sma-3(wk30) III* worms lacking the extrachromosomal array, from the same population, as the experimental comparison group), *crh-1(n3315);sma-3(wk30);wqEx35[Pvha-7∷sma-3 + Pmyo-2∷mCherry]* (with *crh-1(n3315);smo-3(wk30)* worms lacking the extrachromosomal array, from the same population, as the experimental comparison group), *crh-1(n3315);sma-3(wk30);wqEx56[Pdpy-7∷crh-1β∷unc-54 3’ UTR + pY37A1B.5∷crh-1β∷unc-54 3’UTR* + *Pmyo-2∷mCherry]* (with *crh-1(n3315);smo-3(wk30)* worms lacking the extrachromosomal array, from the same population, as the experimental comparison group), *crh-1(n3315);wqEx49[Pdpy-7∷gfp], N2;wqEx49[Pdpy-7∷gfp], N2;wkEx66[Pdpy-7∷wrt-10∷unc-54 3’ UTR* + *PY37A1B.5∷wrt-10∷unc-54 3’ UTR* + *Pmyo-2∷mCherry]* (with N2 worms lacking the extrachromosomal array, from the same population, as the experimental comparison group), *rde-1(mkc36) V;mkcSi13[Psun-1∷rde-1∷sun-1 3’ UTR + unc-119(+)] II;wqex66[Pdpy-7∷wrt-10∷unc-54 3’ UTR + PY37A1B.5∷wrt-10∷unc-54 3’ UTR + Pmyo-2∷mCherry]*. RNAi clones were obtained from the Ahringer RNAi library or were a gift from the Kuwabara lab.

All strains were cultured using standard methods (Brenner, 1974). For experiments, worms were maintained at 20 °C on plates made from nematode growth medium (NGM: 3 g/L NaCl, 2.5 g/L Bacto-peptone, 17 g/L Bacto-agar in distilled water, with 1mL/L cholesterol (5 mg/mL in ethanol), 1 mL/L 1M CaCl_2_,1 mL/L 1M MgSo_4_, and 25 mL/L 1M potassium phosphate buffer (pH 6.0) added to molten agar after autoclaving; Brenner, 1974), while worms collected for hypodermal tissue purification and subsequent RNA sequencing were maintained on high growth medium (HGM: NGM recipe modified as follows: 20 g/L Bacto-peptone, 30 g/L Bacto-agar, and 4mL/L cholesterol (5 mg/mL in ethanol); all other components same as NGM). Plates were seeded with OP50 *E. coli* for *ad libitum* feeding, except for in RNAi experiments. For RNAi experiments, the standard NGM molten agar was supplemented with 1 mL/L 1M IPTG (isopropyl β-d-1-thiogalactopyranoside) and 1mL/L 100mg/mL carbenicillin, and plates were seeded with HT115 *E. coli* containing the RNAi plasmid or empty control vector. To developmentally synchronize experimental animals, hypochlorite-synchronization was performed by collecting eggs from gravid hermaphrodites by exposing them to an alkaline-bleach solution *(e.g.*, 8.0 mL water, 0.5 mL 5N KOH, 1.5 mL sodium hypochlorite), followed by repeated washing of collected eggs in M9 buffer (6 g/L Na_2_HPo_4_, 3 g/L KH_2_Po_4_, 5 g/L NaCl and 1 mL/L 1M MgSo_4_ in distilled water, Brenner, 1974).

### METHOD DETAILS

#### Reproductive span and lifespan assays

As previously described (Luo et al., 2009), hypochlorite-synchronized hermaphrodites were placed as L4 larvae with young (day 1 adult) *fog-2(q71)* males at a 1:3 ratio for 24 h or 48 h of mating, before being individually plated to start the experiment. Mated reproductive spans involved moving individually plated hermaphrodites to fresh plates daily, until progeny production had ceased for two subsequent days. The last day of viable progeny production preceding two days with no progeny was designated as the day of reproductive cessation (assessed by evaluating plates for progeny 1-2 days after the hermaphrodites had been moved to fresh plates). Successful mating was ascertained by evaluating the fraction of male progeny produced. Animals were censored on the day of matricide or loss.

For performing lifespan assays, as previously described (Kenyon et al., 1993), groups of <15 hypochlorite-synchronized hermaphrodites were first placed on individual plates as L4 larvae; they were transferred to freshly seeded plates every two days when producing progeny, and approximately once per week thereafter. Animals were scored as dead when they no longer responded to touch stimuli, and censored on the day of matricide, aberrant vulva structures, or loss/dehydration.

#### Oocyte quality assays

As previously described (Luo et al., 2010; Templeman et al., 2018), hypochlorite-synchronized hermaphrodites were placed as L4 larvae with young (day 1 adult) *fog-2(q71)* males at a 1:3 ratio for 24 h-48 h of mating, before being individually plated; successful mating was ascertained prior to taking images, by evaluating the fraction of male progeny produced in the preceding days. Worms were transferred to freshly seeded plates every 1-2 days thereafter, until oocyte images were taken. For imaging, worms were placed on 4% agarose pads in M9 buffer with 5 mM sodium azide to immobilize them, and differential interference contrast (DIC) microscopy images were captured with a Nikon Eclipse Ti microscope

For scoring oocyte images, the image order was randomized and the scorer was blinded to the genotype represented in each image while assigning a score for each of the signs of deterioration according to the severity of the phenotype. Images were scored based on previously described criteria (Luo et al., 2010; Templeman et al., 2018); in brief, a score of either normal, mild, or severe was assigned for each category, based on the severity of the phenotype. With respect to the ‘cavities’ phenotype, normal indicated no cavities, mild indicated some loss of contact between oocytes, and severe indicated the presence of clear breaks or large gaps between oocytes; with respect to the ‘misshapen oocytes’ category, normal indicated that all oocytes were similarly shaped (generally cuboidal), mild indicated some slight irregularities, and severe indicated one or more oocytes appearing damaged or very irregularly shaped; with respect to the ‘small oocytes’ category, normal indicated no small oocytes, mild indicated that the oocytes did not fill the space between the body wall and germline arm, and severe indicated incidences of very tiny or atrophied oocytes; with respect to the ‘oocytes in uterus’ category, normal indicated no recognizable unfertilized oocytes in the uterus (only embryos), mild indicated at least one recognizable unfertilized oocyte in the uterus, and severe indicated clumped unfertilized oocytes (and likely other cellular material) in the uterus; with respect to the ‘stacked’ category, normal indicated that the developing oocytes of the proximal gonad were essentially all in the same field of view, while severe indicated significant disarray in terms of oocyte organization, with “stacking” of oocytes in multiple fields of view.

#### Late-mating assays

Depending on the experiment outlined in the figure legend, hypochlorite-synchronized eggs were placed on plates seeded with OP50 *E. coli* (for non-RNAi experiments), plates seeded with HT115 *E. coli* containing empty control vector (for “adult-only” RNAi exposure), or plates seeded with HT115 *E. coli* with the RNAi plasmid (for “whole-life” RNAi exposure). Starting on day 1 of adulthood, groups of <25 hermaphrodites were transferred to freshly seeded plates every two days to avoid progeny accumulation; for “adult-only” RNAi experiments, these plates were now seeded with HT115 *E. coli* containing the RNAi plasmid. On the late-mating day in question (usually day 7 of adulthood, unless otherwise stated), individual hermaphrodites were placed on plates seeded with OP50 *E. coli* (for non-RNAi experiments) or HT115 *E. coli* with empty control vector, and three young (day 1 adult) *fog-2(q71)* males were added to each plate. 4-5 days later, plates were assessed for progeny production.

#### Body length and Oil Red O assays

For body size assays, day 1 adult hypochlorite-synchronized hermaphrodites were placed on 4% agarose pads in M9 buffer with 5 mM sodium azide to immobilize them, and DIC microscopy images were captured with a Nikon Eclipse Ti microscope. Body length was assessed using ImageJ v1.5 (Schneider et al., 2012).

For Oil Red O staining (O’Rourke et al., 2009), hypochlorite-synchronized hermaphrodites were collected on day 1 of adulthood and washed with M9 buffer, washed twice with 60% isopropanol, and then exposed to filtered Oil Red O solution overnight (at least 12 h). Worm bodies were then washed with 0.01% Triton X-100 in S-buffer (1.12 g/L K_2_Po_4_, 5.93 g/L KH_2_Po_4_, 5.85 g/L NaCl, pH 6.0), and rinsed several times. To image worms, worms were placed on 4% agarose pads in M9 buffer, and color images were captured with a Nikon Eclipse Ti microscope. Worm images were analyzed for mean intensity using CellProfiler v3.0 (Wählby et al., 2012).

#### FACS isolation of dissociated hypodermal tissue cells

Hypodermal tissue was collected for FACS isolation as previously described (Kaletsky et al., 2016; 2018). In brief, day 1 adult hypochlorite-synchronized hermaphrodites were collected from HGM plates and washed several times with M9 buffer to remove excess bacteria. The pellet was washed with and then resuspended in lysis buffer (200 mM DTT, 0.25% SDS, 20 mM HEPES pH 8.0, 3% sucrose); worms were incubated in lysis buffer for 6.5 min at room temperature. The pellet was rapidly washed 5 times with M9 buffer, and resuspended in freshly prepared 20 mg/mL pronase from *Streptomyces griseus* (Sigma-Aldrich) in Leibovitz’s L-15 buffer (ThermoFisher Scientific; adjusted to 340 mM osmolarity with sucrose). Resuspended worm bodies were incubated at room temperature for <20 min with periodic mechanical disruption by frequent vigorous pipetting. When most worm bodies had been dissociated, ice-cold L-15 buffer containing 2% fetal bovine serum (FBS; Gibco, ThermoFisher Scientific) was added, and samples were passed over a pre-wet 20 mm nylon filter (Sefar Filtration) using gravity flow.

Filtered cell material was diluted in ice-cold L-15 buffer/2% FBS and kept on ice, before being sorted shortly thereafter using a BD FACSAria Fusion (BD Biosciences; 70 μm nozzle, 488 nm laser excitation). Gates for GFP detection were set by comparing to wild-type (non-fluorescent) cell suspensions prepared in parallel from a population of non-fluorescent N2 worms that was hypochlorite-synchronized and maintained alongside the experimental samples. Positive fluorescent events were sorted directly into Eppendorf tubes containing TRIzol reagent (ThermoFisher Scientific) for storage at −80 °C, prior to RNA extraction. For each sample, approximately 80,000-250,000 GFP-positive events were collected.

#### RNA isolation, amplification, library preparation, and sequencing

As previously described (Kaletsky et al., 2016; 2018), RNA was extracted using a standard TRIzol/chloroform/isopropanol method, and cleaned using Qiagen RNEasy MinElute kit, including the DNase digestion step (Qiagen), following the manufacturer’s instructions. Total RNA quality and quantity were assessed with an Agilent Bioanalyzer. The SMARTer Stranded Total RNA-Seq v2 - Pico Input Mammalian kit (Takara Bio) was used for preparation, amplification, and purification of Illumina sequencing libraries (skipping the rRNA depletion steps), as per manufacturer-suggested practices. Libraries were quantified with an Agilent Bioanalyzer, and then submitted for sequencing on the Illumina HiSeq 2000 platform.

#### RNA sequencing data analysis

RNA sequencing analysis was performed as previously described (Kaletsky et al., 2016; 2018). In brief, FASTQC was used to inspect the quality scores of the raw sequence data and adapter-cut/trimmed data, and look for biases. The first 5 bases of each read were trimmed before adapter trimming to remove the universal Illumina adapter and impose a base quality score cutoff of 20, using Cutadapt v1.6. The trimmed reads were mapped to the *C. elegans* genome (Ensembl 84/WormBase 235) using STAR (Dobin et al., 2013) with Ensembl gene model annotations, using default parameters for single-end reads. The reads aligning to each gene were counted with HTSeq, and DESeq2 (Love et al., 2014) was used for differential expression analysis. Genes with an adjusted p-value ≤ 0.05 were considered to be significantly differentially expressed. g:Profiler (Reimand et al., 2007) was used for GO term analysis, and GO terms with an adjusted p-value ≤ 0.05 were considered to be significant.

#### Gene Set Enrichment Analysis

Gene Set Enrichment Analysis (GSEA) software (Subramanian et al., 2005), which identifies statistically significant concordant changes in comparisons of two genotypes, was used to test whether the whole-body transcriptional profile of *crh-1(-)* mutants versus wild type (Lakhina et al., 2015) exhibits significant positive enrichment for the up- and down-regulated genes identified in the whole-body transcriptional profile *sma-2(-)* TGF-β Sma/Mab signaling mutant L4 larvae versus wild type (Luo et al., 2010). All data sets were previously published.

#### Images and visualization

RStudio v1.2 was used to create the volcano plot. Visualization of overlap between gene lists was generated using BioVenn (Hulsen et al., 2008). Some components of drawn models were created using BioRender.

### QUANTIFICATION AND STATISTICAL ANALYSIS

Lifespan and reproductive span assays were assessed using standard Kaplan-Meier log rank survival tests, with the first day of adulthood of synchronized hermaphrodites defined as t = 0. For oocyte quality experiments, chi square analyses were used to determine whether there were significant differences between populations for each category of scored oocyte phenotypes. For late-mating assays, chi square tests were used to compare the percentages of worm populations capable of progeny production, with Bonferroni corrections applied to adjust for multiple comparisons. Unpaired t tests were used for body length or Oil Red O staining intensity comparisons between genotypes. All experiments were repeated on separate days with separate, independent populations, to confirm that results were reproducible. Prism 8 software was used for statistical analyses; software and further statistical details used for RNA sequencing analyses are described in the method details section of the STAR methods. Additional statistical details of experiments, including sample size (with n representing the number of worms), can be found in the figure legends.

## DATA AND SOFTWARE AVAILABILITY

RNA sequencing data will be publicly deposited.

## Supplemental Information Titles and Legends

**Figure S1. Loss of CREB reduces growth and lipid storage**. *Related to Figure 1.*

(A) Reduced body length in day 1 adult *crh-1(n3315)* (n = 32) versus N2 (n = 36).

(B) Reduced whole-body lipid storage, as indicated by Oil Red O staining intensity, in day 1 adult *crh-1(n3315)* (n = 25) versus N2 (n = 20).

Boxes represent 25th to 75th percentiles, with lines at the means, points represent individual worms, whiskers represent minimum and maximum values. **** *p* ≤ 0.0001.

**Figure S2. Similarities between CREB and TGF-β Sma/Mab signaling**. *Related to Figure 2.*

(A) *crh-1(-)* mutants have similar transcriptional profiles to *sma-2(-)* mutants. (Left) Up-regulated genes in *sma-2(e502)* versus N2 (Luo et al., 2010) are positively enriched in *crh-1(n3315)* mutants versus N2 (Lakhina et al., 2015). Normalized Enrichment Score 2.20; nominal *p*-value 0.0; FDR q-value 0.0. (Right) Down-regulated genes in *sma-2(e502)* versus N2 (Luo et al., 2010) are positively enriched in *crh-1(n3315)* mutants versus N2 (Lakhina et al., 2015). Normalized Enrichment Score −1.77; nominal *p*-value 0.0; FDR q-value 0.0. The x-axes show a heatmap of gene clustering, while the y-axes display gene enrichment scores compared to *sma-2(-)* transcriptional changes.

(B) Hypodermal *crh-1* rescue is sufficient to increase growth of small *crh-1(-)* mutants Hypodermal *crh-1* rescue *(crh-1*(-);hypo CREB, *i.e., crh-1(n3315);Pdpy-7∷crh-1β* + *PY37A1B.5∷crh-1β*, n = 24) significantly increases the body lengths of *crh-1(n3315)* worms (n = 28). Boxes represent 25th to 75th percentiles, with lines at the means, points represent individual worms, whiskers represent minimum and maximum values. ** *p* ≤ 0.01.

**Figure S3. Altered genes in both adult CREB hypodermal tissue and *sma-2(-)* TGF-β Sma/Mab mutant larval whole bodies.** *Related to Figure 5.*

(A and B) Venn diagrams showing overlap between the sets of genes that were significantly (A) upregulated or (B) downregulated in hypodermal tissue of day 1 adult *crh-1(n3315)* mutants versus N2 (refer to Figure 4, and Table S1), and the sets of genes that were significantly (A) upregulated or (B) downregulated in the whole bodies of *sma-2(e502)* TGF-β Sma/Mab signaling mutant L4 larvae (Luo et al., 2010).

**Table S1. Genes significantly up- or down-regulated in *crh-1(n3315)* versus N2 hypodermal tissue.** *Related to Figure 4.*

*See separate excel files for content*

**Table S2. GO terms significantly enriched for genes up- or down-regulated in *crh-1(n3315)* versus N2 hypodermal tissue.** *Related to Figure 4.*

*See separate excel files for content*

