## Supplementary figures and images for "CREB non-autonomously regulates Reproductive Aging through Hedgehog/Patched Signalling"

### Figure S1

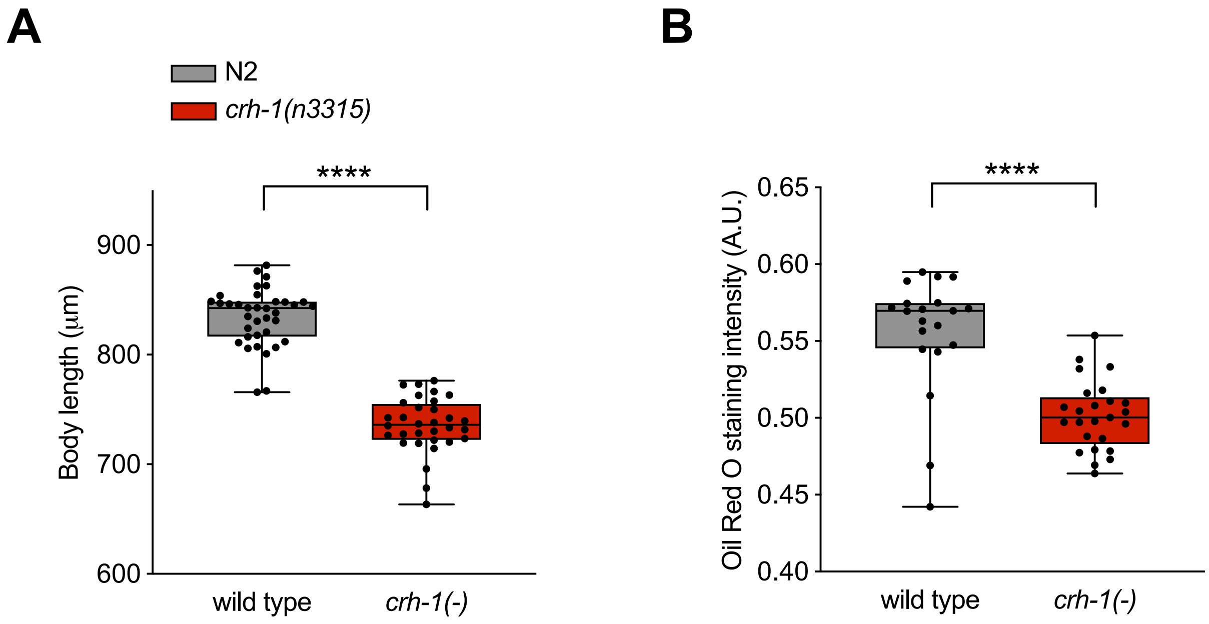

### Figure S2

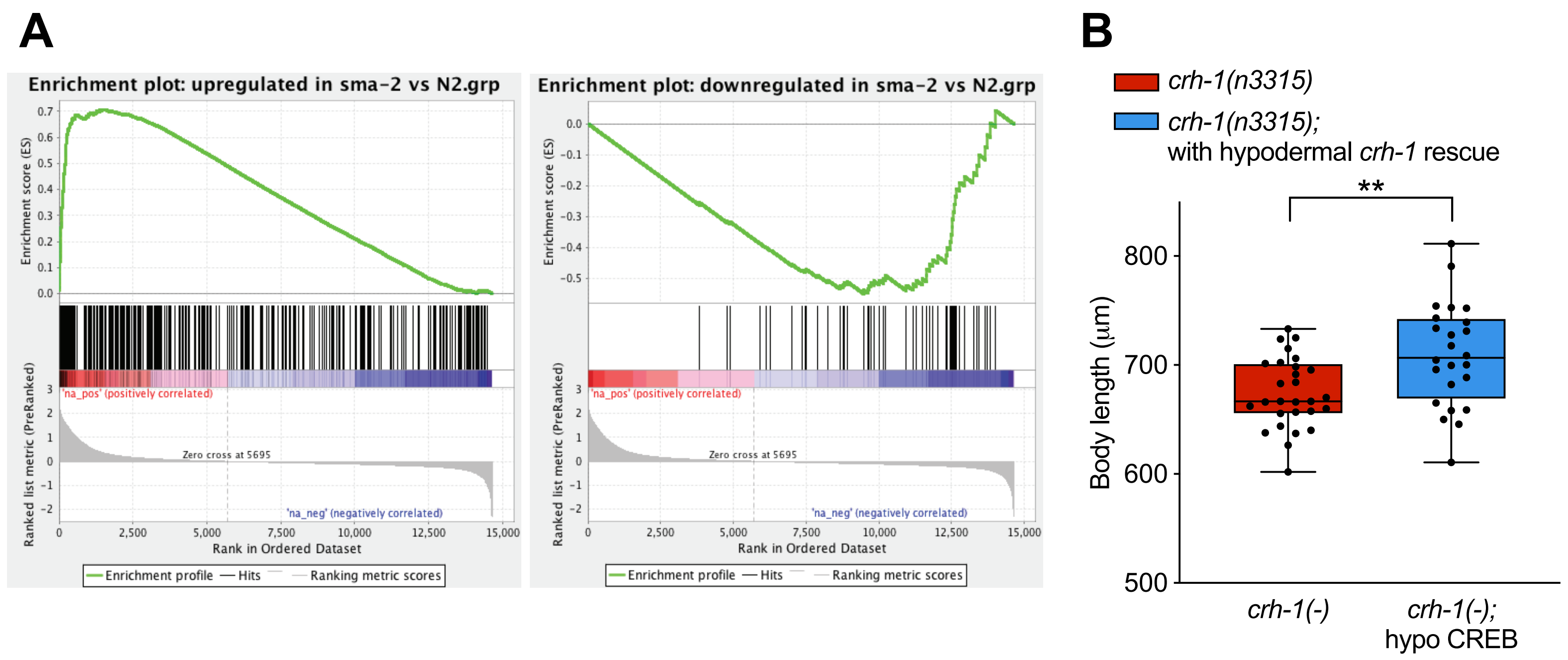

### Figure S3

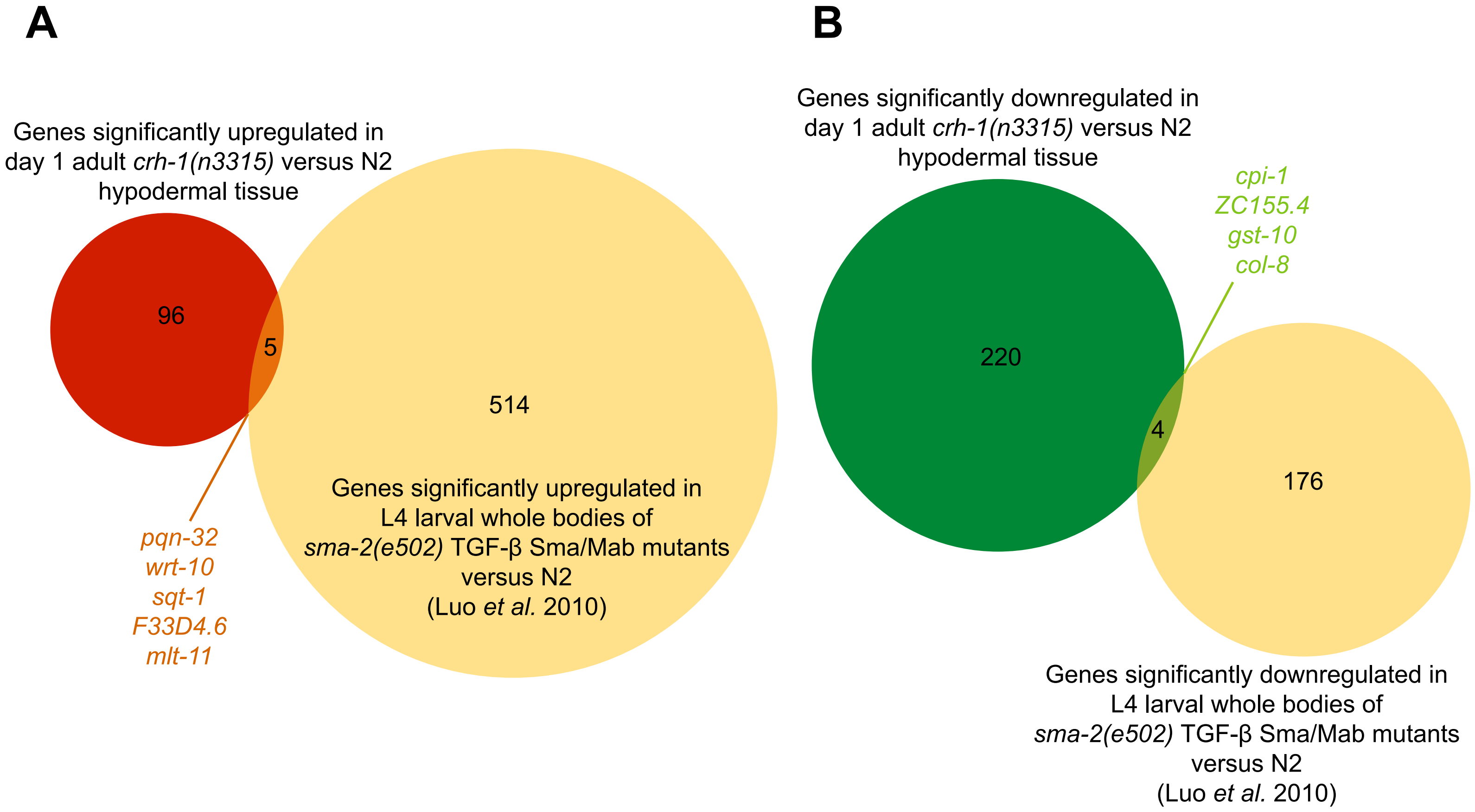
